# Enhanced metagenomics-enabled transmission inference with TRACS

**DOI:** 10.1101/2024.08.19.608527

**Authors:** Gerry Tonkin-Hill, Yan Shao, Alexander E. Zarebski, Sudaraka Mallawaarachchi, Ouli Xie, Tommi Mäklin, Harry A. Thorpe, Mark R. Davies, Stephen D. Bentley, Trevor D. Lawley, Jukka Corander

## Abstract

Coexisting strains of the same species within the human microbiota pose a substantial challenge to inferring the host-to-host transmission of both pathogenic and commensal microbes. Here, we present TRACS, a highly accurate algorithm for estimating genetic distances between strains at the level of individual SNPs, which is robust to intra-species diversity within the host. Analysis of well-characterised Faecal Microbiota Transplantation datasets, along with extensive simulations, demonstrates that TRACS substantially outperforms existing strain aware transmission inference methods. We use TRACS to infer transmission networks in patients colonised with multiple strains, including SARS-CoV-2 amplicon sequencing data from UK hospitals, deep population sequencing data of *Streptococcus pneumoniae* and single-cell genome sequencing data from malaria patients infected with *Plasmodium falciparum*. Applying TRACS to gut metagenomic samples from a large cohort of 176 mothers and 1,288 infants born in UK hospitals revealed species-specific transmission rates between mothers and their infants. Notably, TRACS identified increased persistence of *Bifidobacterium* breve in infants, a finding missed by previous analyses due to the presence of multiple strains.

## Introduction

Host-to-host transmission is a fundamental process that shapes the interactions between humans and microbes. Tracking the spread of pathogens using genomics has become a major tool in public health, helping to prevent the spread of disease at both local and global scales(1, 2). Beyond pathogens, understanding the transmission and colonisation dynamics of commensal microbes would greatly improve our understanding of microbiome assembly and maintenance, and how this is influenced by diet, lifestyle, culture, clinical interventions and social interactions. In addition, the human microbiome contains microbes with therapeutic potential to treat a variety of human diseases. The ability to identify candidate strains during clinical interventions such as Faecal Microbiota Transplantation (FMTs) and Live Biotherapeutic products would greatly de-risk and accelerate the development of microbiome-based therapeutics.

Whole genome sequencing (WGS) has reshaped our ability to infer transmission chains by enabling the detection of single-nucleotide variants, facilitating precise tracking of slowly-evolving pathogens such as Methicillin-resistance *Staphylococcus aureus* (MRSA)(2). However, current WGS approaches often focus on a representative strain genome of a single species, overlooking the microbial strain diversity within individuals(3). Recent efforts have begun to address this limitation using metagenomics, enabling the simultaneous consideration of multiple species and their strains in a single analysis (4, 5). Deep population sequencing of specific species can be achieved through targeted enrichment techniques like culture or PCR amplicon sequencing, allowing for fine-scale analyses of multiple co-colonising strains (6–8). Existing gold-standard methods for tracking transmission using metagenomic or deep population sequencing data have typically been developed for curated academic studies (5, 9–11). While these methods offer valuable insights, such as within-host variant calling and identification of selection pressures via dN/dS, they lack the speed and flexibility required for routine public health monitoring. Notably, they often lack the temporal resolution needed to accurately distinguish between strains transmitted recently (weeks to months) and distantly related genomes (years).

Tools that rely on reference marker gene databases including MIDAS (11, 12) and StrainPhlAn (9) consider only a small portion of a species genome (10-200 genes(4)) and do not attempt to separate within species diversity. This substantially limits the temporal resolution at which transmission can be inferred. An alternative approach, used by StrainGE (10) and inStrain (database mode)(5) involves identifying the species found across the entire dataset and building a dataset-specific reference genome database for read alignment. This approach relies heavily on the similarity between the transmitted and reference genome and does not allow for the continuous integration of new samples into an analysis, making it unsuitable for routine genomic surveillance.

Another common approach relies on de novo metagenomic assembly, including the inStrain (assembly mode)(5) and STRONG(13) pipelines. Assembly requires high sequencing coverage of the transmitted genome(10). These methods generally perform best when samples are pooled before assembly, followed by genome binning in a co-assembly workflow. However, to avoid extensive genome deduplication, which effectively turns the process into a reference-based approach, they must be applied to pairs or reduced subsets of samples. This can substantially increase the computational burden. Recombination and shared homology between strains of the same species within a sample is also generally not accounted for by existing algorithms, which can have a major impact on the accuracy of transmission inference when considering metagenomic and population sequencing data. To address these issues, we developed TRAnsmision Clustering of Strains (TRACS), a highly accurate and easy-to-use algorithm for establishing whether two samples are plausibly related by a recent transmission event.

The TRACS algorithm distinguishes the transmission of closely related strains by identifying genetic differences as small as a few Single Nucleotide Polymorphisms (SNPs), which is crucial when considering slow-evolving pathogens. The algorithm employs novel statistical filtering techniques to account for variable sequence coverage, shared homology between strains, and sequencing errors. Critically, TRACS was designed to estimate an accurate but conservative (*lower bound*) of the SNP distance and considers each reference alignment independently, enabling continuous integration of new samples. This algorithmic approach allows TRACS to function similarly to how SNP distances are used in conventional single-isolate genomic epidemiology studies, making it an ideal tool for accurately identifying putative transmission networks and ruling out transmission events in ongoing public health applications. However, similar to single-isolate studies, discerning fine-scale transmission structure, such as the direction of transmission, typically requires additional epidemiological data, such as contact tracing information.

We demonstrate using comprehensive simulations, and by considering well characterised Faecal Microbiota Transplant (FMT) metagenomic datasets that TRACS has superior performance to existing metagenomic transmission inference methods and can reliably identify plausible transmission events. We apply TRACS to a diverse set of pathogens including viruses, bacteria and parasites, demonstrating the scalability and versatility of the algorithm. Finally we consider the development of the infant gut microbiome and species specific transmission rates between mothers and infants using a large cohort of 1,464 mothers and infants.

## Results

### TRACS addresses key challenges in metagenomic transmission inference

The TRACS algorithm reliably estimates pairwise genetic SNP distances between genomes in complex metagenomic or deep population sequencing data. The algorithm allows for the efficient addition of new samples to an analysis, a key requirement for near real-time use in public health settings, such as emerging outbreaks.

The initial ‘alignment’ stage of TRACS uses the hash-based search algorithm ‘Sourmash’ to identify a set of reference genomes that best represent the species and their strains in a sample(14). Unlike alternative approaches, TRACS avoids the error prone deconvolution of reads into different genome bins and instead aligns the entire read set to each reference genome individually, generating a count of the alleles observed at each site(15). This enables the incremental addition of new samples and the integration of novel species into the reference database without requiring the reprocessing existing data.

A novel set of statistical filtering algorithms are then applied to exclude regions affected by shared sequence homology, multi-mapping reads, poor alignment quality, and low sequencing coverage (Figure 1). This includes a scan statistic, similar to those used in phylogenetics to detect recombination(16), that can identify regions with elevated polymorphism rates, often resulting from shared sequence homology between co-colonising strains or gene duplications. TRACS also includes an empirical Bayes approach to account for regions of a reference genome with insufficient coverage to accurately represent multiple strains present within a sample (described in Methods). The resulting filtered alignments are converted into reference based Multiple Sequence Alignments (MSAs) for each reference genome. Finally, TRACS incorporates a fast, IUPAC-aware pairwise SNP distance estimation algorithm, enabling rapid inference of genetic distances.

**Fig. 1.**
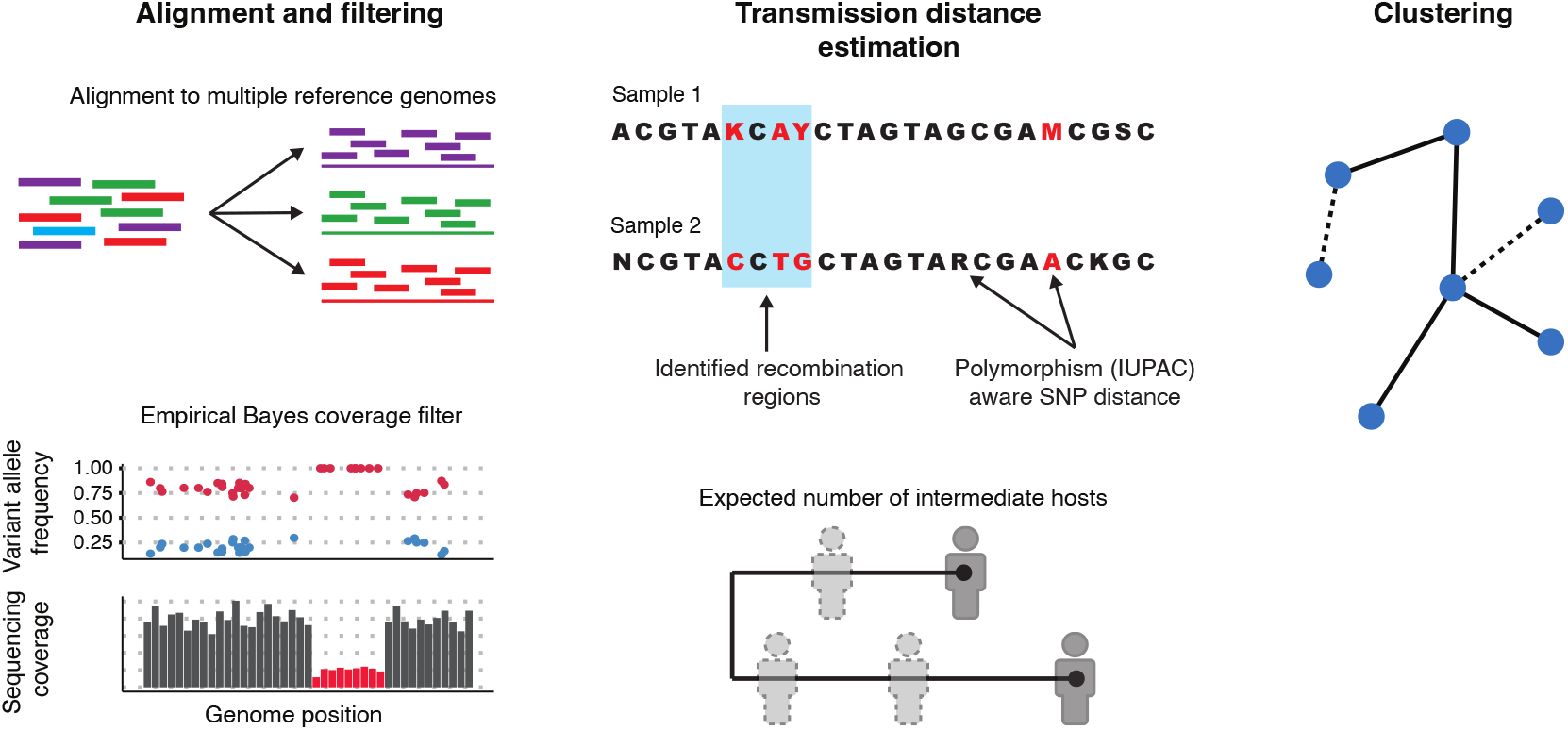
A schematic illustrating the key components of the TRACS algorithm. Left: reads are aligned to each reference genome identified by Sourmash separately. An empirical Bayes method is used to pinpoint genome regions with insufficient coverage for minority strain identification. Centre: alignments from each sample are transformed into a multiple sequence alignment using IUPAC ambiguity codes to represent multiple alleles at a single site. Rapid pairwise SNP distances are then calculated, excluding potential recombination regions by identifying areas with high SNP density. The TransCluster algorithm can optionally be applied to estimate the expected number of intermediate hosts between two samples. Right: the resulting pairwise transmission distance estimates are clustered using single linkage hierarchical clustering to infer putative transmission clusters. Transmission distance thresholds are inferred using a mixture distribution to separate sample pairs that are known to be distantly related from those that include recent transmissions.

Optionally, TRACS can incorporate sampling dates and a known transmission generation time to estimate the expected number of intermediate hosts separating two samples. In addition to improved computational speed, this enhances the interpretability of the TransCluster algorithm (17), which only estimated the probability of a given number of intermediate hosts (Methods). While the alignment module of TRACS is designed for metagenomic and population sequencing data, MSAs produced by alternative tools designed for isolate-data, such as Snippy(18), can also be used directly as input to the TRACS distance module.

A key challenge in using pairwise distance methods to infer transmission chains is determining an appropriate threshold for when two strains are likely connected by recent transmission. TRACS introduces a novel method that uses a mixture distribution to distinguish between recent transmission events and strains that are distantly related by years of evolution. This results in thresholds that are more specific than those provided by previous methods such as those that rely on Youden’s index (19). Similar to previous efforts (4), this approach leverages data from known closely related samples, such as strains from the same individual, and distinguishes them from distantly related samples.

The TRACS algorithm is implemented in python and C++, and is available under the MIT open source licence at https://github.com/gtonkinhill/tracs/.

### TRACS achieves superior performance on simulated transmission pairs

To assess the ability of TRACS to accurately estimate small genetic distances in pairs of samples containing multiple strains of the same species, we simulated WGS reads from mixtures of common *Streptococcus pneumoniae* strains. Common transmission SNP thresholds for *S. pneumoniae* are often *<* 10 SNPs(20). In each pair we selected one genome to be shared between samples at a specified SNP distance (5, 50 and 500 SNPs) ensuring a minimum average sequencing coverage of 5x for the transmitted strain. The TRACS algorithm was compared to InStrain, StrainGE, and StrainPhlAn.

All algorithms except TRACS substantially overestimated genetic distances (Figure 2A). This bias was larger among low-frequency strains (Supplementary Figure 1), limiting the ability of these algorithms to reliably rule out recent transmission events, as commonly used SNP distance thresholds for many species fall well below the resulting inflated estimates (20). StrainGE was the best-performing algorithm after TRACS, likely due to its ability to account for minority strains and its use of competitive mapping among genomes within a species. This approach helps mitigate the effects of horizontal gene transfer (HGT) and the resulting shared homology between strains, as demonstrated in the original StrainGE publication(10). However, StrainGE only reported relatively accurate SNP distances at higher values (500 SNPs), which substantially exceeds common thresholds (typically <50 SNPs) used to distinguish transmission in *S. pneumoniae* (21). To determine which filter in the TRACS algorithm had the greatest impact, we evaluated combinations of the novel statistical filtering methods (Supplementary Figure 2). The coverage filter, including the Empirical Bayes algorithm, produced the largest improvement, followed by the recombination filter. Repeating these simulations using an in silico model of high-accuracy Nanopore R10.4.1 reads further demonstrated TRACS’ compatibility with multiple high-accuracy sequencing technologies (Supplementary Figure 5).

**Fig. 2.**
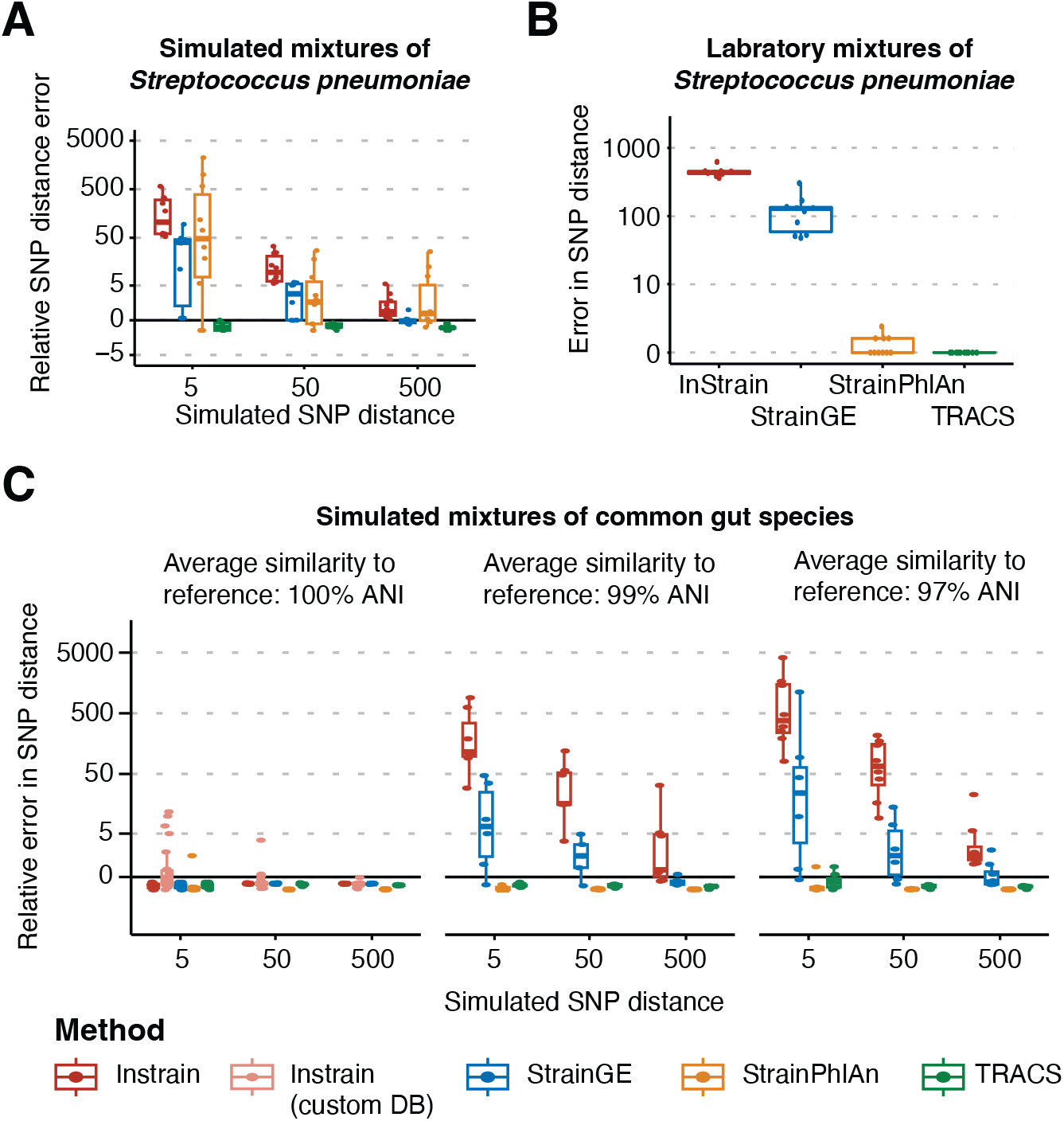
Accuracy of SNP distance estimation algorithms on simulated and laboratory mixtures. A) The relative error (*d*_estimated_ − *d*_simulated_)*/d*_simulated_, in the estimated pairwise SNP distance across four algorithms, when applied to a simulated mixture of pneumococcal genomes. A single genome is simulated to have been transmitted between each pair of samples with a SNP distance given on the x-axis. Ten replicate simulations were performed for each parameter set with boxplots indicating the distribution in relative error rates. Points closest to zero are the best performing, with positive values indicating an overestimation of the SNP distance and negative values indicating an underestimation. B) The results of running each algorithm on artificial laboratory mixtures of pneumococcal strains from Knight et al.(22). Each pair of samples included at least one identical strain and thus the correct SNP distance should be zero. Except for TRACS, all algorithms erroneously estimate elevated SNP distances. C) Simulated transmission of individual species was modelled between synthetic gut metagenome samples. Genomes were selected to have average nucleotide identities (ANI) of 100%, 99%, and 97% relative to the GTDB representative genomes, which serve as the reference database for the InStrain, StrainGE, and TRACS algorithms. StrainPhlan was run using the marker gene database (v4.0.5). Similar to the pneumococcal simulation, each of the ten species was simulated independently at three different SNP distance thresholds: 5, 50, and 500 SNPs. Only a single strain per species was included in each simulation. In addition to running InStrain in reference-based mode using GTDB representative genomes, InStrain was also run with custom reference genomes (custom DB) generated for each sample pair using metaSPAdes v4.2.0. Contigs were ‘binned’ into species by aligning the assemblies back to the simulated genomes.

To validate these results with real sequencing data, with a known ground truth, we analysed 13 laboratory mixtures of different *S. pneumoniae* strains from a previous study (22). In each case, at least one identical pneumococcal genome was shared between samples, meaning all algorithms should infer zero SNPs. As shown in Figure 2B, TRACS was the only algorithm that reliably inferred zero SNPs in all cases. Strain-PhlAn correctly identified zero SNPs in some cases; however, its reliance on marker genes can lead it to systematically underestimate SNP distances, as shown in the subsequent analysis of FMT data. To further explore the reconstruction of transmission chains from metagenomic data involving multiple distinct species, we simulated genome mixtures representing common gut bacteria at varying levels of similarity to the reference database, 100%, 99%, and 97% pairwise sequence identity (see Methods). In these simulations, a single species identified in a previous study of transmission using metagenomics was simulated to transmit with a specified SNP distance (23). A common reference database was used for all methods with the exception of StrainPhlAn where the default marker gene database v4.0.5 was used. TRACS remained the best performing algorithm under these settings (Figure 2C). InStrain consistently overestimated SNP distances for most simulated pairs once the transmitted genome diverged by at least 1% from the reference database (Figure 2C). Alternative strategies that generate study-specific reference databases, such as co-assembly of all samples or individual sample assembly followed by deduplication, could potentially improve these estimates. However, because inStrain relies on competitive read mapping, it is recommended that its reference databases be deduplicated so that any pair of genomes shares no more than 98% sequence identity. Consequently, in studies where multiple strains of the same species are present (a common occurrence in large datasets), strains that diverge from the assembled and/or deduplicated reference genome would still produce errors similar to those shown in Figure 2C.

As an alternative, we explored generating sample-pair–specific reference genomes. In this mode (“inStrain custom DB”), metaSPAdes was used to assemble each pair of samples independently, producing a unique reference database for each pair (Figure 2C). While this approach improved SNP distance estimates, its computational cost is prohibitive for larger studies. Excluding the computational cost of assembly, using pair-specific references requires the inStrain alignment and variant-calling steps to be run separately for each pair. This step alone required approximately 12.5 CPU hours per pair (Supplementary Figure 7). For a modest dataset of 100 samples, this would translate to 4,950 separate alignment steps and more than 2.5k CPU hours. Furthermore, if multiple strains of the same species were present within a given pair, this approach might still fail to assemble the transmitted strain as the reference genome.

### Enhanced estimates of pathogen transmission across diverse taxonomic groups in high-burden settings

A major advantage of the TRACS algorithm is its ability to identify putative transmission events in cases where the hosts are colonised with multiple strains of the same species. This is a major concern in many high disease burden settings. Unlike alternative algorithms such as StrainPhlAn, the TRACS algorithm is applicable across diverse taxonomic groups including parasitic, viral and bacterial species. To demonstrate the effectiveness of the TRACS algorithm across a wide range of pathogens we considered well characterised datasets across three different settings.

#### SARS-CoV-2

Although infection with multiple SARS-CoV-2 strains is relatively rare (24), the rates of occurrence can be higher in locations with high disease burdens, such as within hospital wards, particularly when infection control procedures have broken down.

To demonstrate the ability of the TRACS algorithm to account for this challenges, we considered 37 of 1181 SARS-CoV-2 samples, collected from the East of England in early 2020 which were processed using Illumina deep amplicon sequencing(25). These samples were found to contain multiple distinct strains after sequencing each sample in replicate to account for sequencing errors (26). These were compared to samples containing a single strain within the mixture. Thus, the smallest SNP distance between samples should be zero. To investigate the utility of applying TRACS to these samples, we compared the inferred SNP distance using TRACS to a consensus based approach, with and without filtering for problematic sites such as hypermutable loci and regions impacted by the ends of amplicon sequencing reads (Figure 3A) (27).

**Fig. 3.**
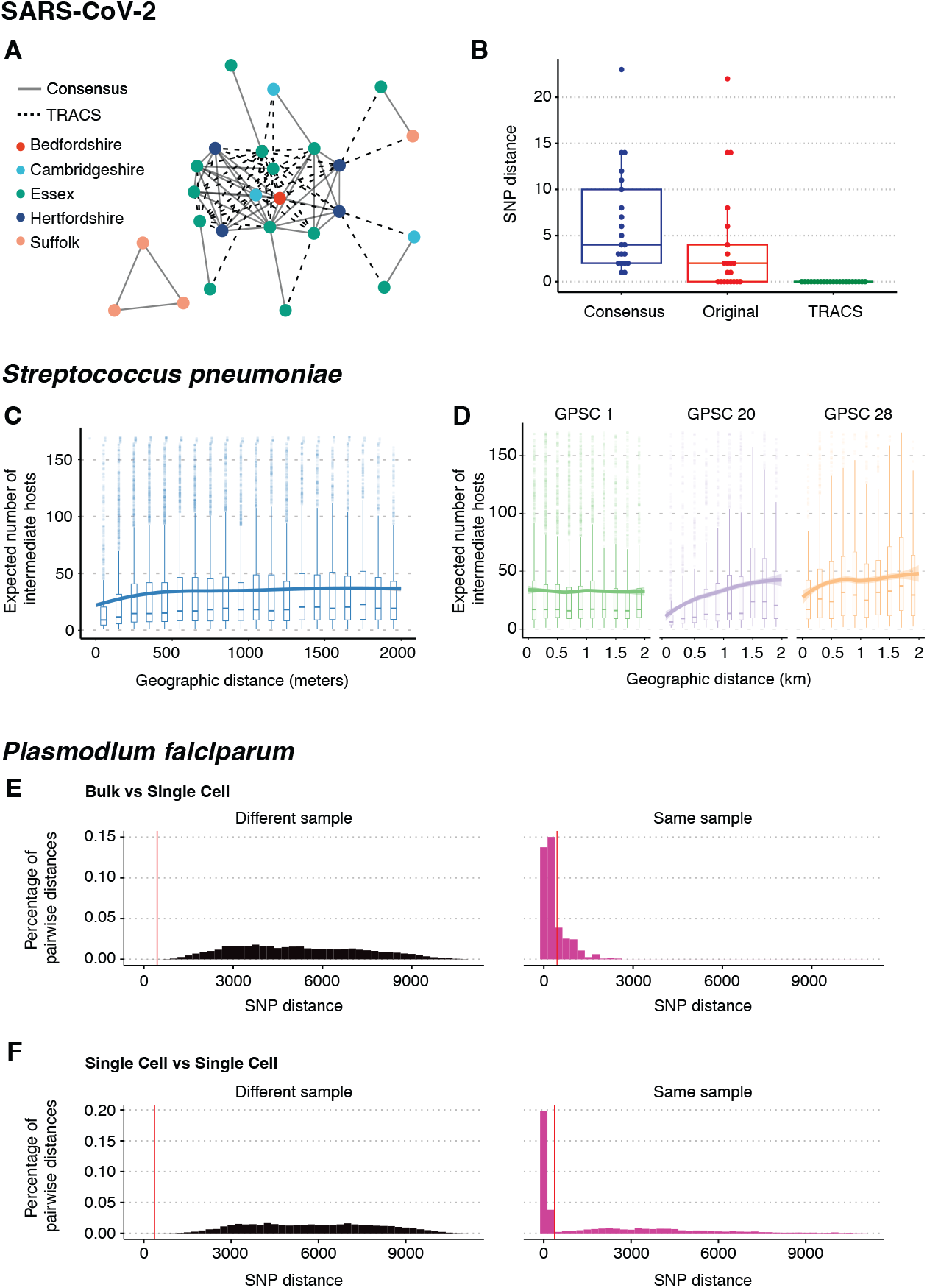
Transmission inference across diverse taxonomic groups using TRACS. A) A SARS-CoV-2 transmission network comprising samples that contain multiple distinct strains. Solid lines indicate transmission events observable using both TRACS and the typical consensus-based sequencing, while dotted lines represent additional links identified by TRACS alone. B) The inferred SNP distance between pairs of samples containing the same strain, where at least one sample includes multiple distinct strains. The consensus approach considers the dominant allele across the entire SARS-CoV-2 reference genome, whereas the ‘original’ approach excludes sites frequently filtered out to avoid hypervariable or error-prone regions. The TRACS algorithm correctly infers zero SNPs in all pairs. Boxplots indicate the distribution of the inferred number of SNPS. The central line represents the median, the edges of the box indicate the interquartile range (IQR), and the whiskers extend to 1.5 times the IQR. C) The expected number of intermediate hosts between each pair of pneumococcal samples taken from different individuals in the Maela refugee camp versus the geographic distance between their homes within the camp with a LOESS smoothing. The shaded area indicates the corresponding confidence interval. Boxplots show the median expected number of intermediate hosts. The edges of each box indicate the interquartile range (IQR), and the whiskers extend to 1.5 times the IQR. Sample pairs with a divergence time outside the establishment of the camp in 1984 were excluded. D) Similar to C, but only points involved in the three most common global pneumococcal sequence clusters (GPSCs) are shown. A strong geographic signal is absent for the non multi-drug resistant lineage GPSC 1, which is known to have a longer carriage duration. E) and F.) The distribution of SNP distances inferred by TRACS for single cell vs. single cell, and bulk vs. single cell samples. The vertical red line indicates the SNP threshold inferred using the TRACS mixture distribution approach. The high number of short SNP distances between single cells from the same sample indicates that TRACS can accurately distinguish closely related genomes within mixed infections of *P. falciparum*.

When the shared strain was in the minority, a consensus based approach consistently overestimated the SNP distance between samples despite filtering out problematic regions. In contrast, TRACS correctly inferred 0 SNPs between samples in all cases without requiring manual filtering of problematic sites. Figure 3B indicates the transmission network inferred assuming 0 SNPs between strains for those mixed samples with a geographic location. The dotted edges indicate transmission links that would have been missed using a standard consensus based framework. This example highlights the ability of TRACS to both account for the presence of multiple strains and to robustly control for the impacts of sequencing errors and hypermutable sites that frequently result in polymorphic variants within samples. TRACS is also likely to be robust to rare cases of recombination between strains within the host. In this case, ignoring polymorphisms could cause consensus methods to overestimate SNP distances, whereas, with sufficient coverage of both parent strains, TRACS would still detect shared strains between samples.

#### Streptococcus pneumoniae

An added benefit of the TRACS algorithm is its ability to estimate the expected number of intermediate hosts between two samples by incorporating sampling dates. Dates can be used as an additional piece of information to rule out transmission for species or lineages with low sequence diversity (17).

To demonstrate this approach, we considered 3,761 nasopharyngeal swabs taken from 468 infants and 145 of their mothers living in a refugee camp in Thailand, where culture based enrichment and whole pneumococcal population Illumina deep sequencing had been performed as part of a previous study (7). TRACS was run on this dataset using a custom database of pneumococcal reference genomes from the Global Pneumococcal Sequencing project (28). Consistent with previous studies, we assume a molecular clock rate of 5.3 SNPs/year and a transmission generation time of two months (7). After running TRACS, the expected number of intermediate hosts between all pairs was compared to the geographic distance separating the households of participants (Figure 3C-D). Although the original TransCluster algorithm was previously applied to this dataset, it only estimates individual probabilities, such as the probability of direct transmission, without estimating the overall expected number of intermediate hosts.

There was a strong correlation between geographic distance and the expected number of intermediate hosts, despite the small size of the refugee camp (2.4 km^2^). Interestingly, this correlation was not found for all the major lineages. In particular, common multi-drug resistant lineages such as Global Pneumococcal Sequencing Cluster (GPSC) 1, had no clear association between transmission and geographic distance within the camp (Figure 3D). These lineages are known to have longer carriage durations, which can obscure geographic transmission signals (29, 30). Thus the observed signals are likely driven by lineages that transmit faster and exhibit shorter carriage durations, such as GPSC 20. Understanding the different dynamics governing the transmission of these lineages is crucial for the design of interventions aimed at reducing pneumococcal disease.

#### Plasmodium falciparum

The increased genome size of major parasitic pathogens has hampered the adoption of routine WGS in disease surveillance. However, the rapidly decreasing cost of sequencing is leading to the increasing use of WGS to track major parasite populations including the malaria parasite, *P. falciparum*, which causes over half a million deaths each year (31). Multiple strains of *P. falciparum* are frequently found within people living in endemic areas with high burdens of disease (32). To investigate the ability of the TRACS algorithm to accurately identify shared strains within mixed *P. falciparum* infections, we considered a dataset involving 49 samples from Chikh-wawa, Malawi that were positive for *P. falciparum* (33). Both Illumina bulk sequencing of mixed populations in addition to single cell and single clone enrichment had been performed as part of a previous study leading to 49 mixed WGS samples and 509 single genomes(33). Although the sparse sampling makes direct transmission between samples unlikely, the combination of bulk and single-cell sequencing allows us to assess whether the TRACS algorithm can accurately detect shared strains within these mixed samples. Figures 3E and 3F indicate the distribution of the inferred SNP distances between single cell genomes and bulk samples from either the same sample or between samples. The clear separation of SNP distances between genomes sequenced from the same sample and between samples indicates that TRACS can accurately identify shared strains within this data. As WGS of malaria parasites becomes more routine in public health settings, the TRACS algorithm can be used to accurately determine if a pair of samples could be related by a recent transmission event. A fine scale understanding of transmission dynamics in high burden settings will aid in the design of strategies aimed at reducing and eventually eliminating malaria.

### Improved estimates of engraftment in faecal microbiota transplant triads

Faecal microbiota transplantation (FMT) involves transferring a donor stool sample to a recipient, often via orally-delivered capsules, and has demonstrated patient benefit to treat infections, autoimmune diseases, graft versus host diseases and cancers. Development of microbiome drugs often requires the identification of specific bacterial strains with beneficial properties from complex microbiomes containing 100-1000s strains. To further evaluate the efficacy of the TRACS algorithm in metagenomics applications, we analysed an extensively characterised Faecal Microbiota Transplantation (FMT) dataset with well-defined transmission (or engraftment) relationships between samples(34). The capability of each algorithm to detect engraftment links between human gut microbiome samples from donors and recipients was examined by considering 23 previously published FMT triads(34). Metagenomic samples were originally collected from donors and recipients before and after transplantation across three patient cohorts, including individuals with *Clostridium difficile* infections, inflammatory bowel disease (IBD), and recurrent multi-drug resistant infections.

The TRACS algorithm was run on each of these samples and the results compared with StrainPhlAn, InStrain and StrainGE (Figure 4A-C, Supplementary Figure 4). As StrainGE requires substantial computing resources to consider transmission of every observed genus we restricted this analysis to *Bifidobacteria*. The inferred number of engraftment events was then inferred, comparing samples from the same patient post-FMT and those from different cohorts (Figure 4A). Species specific SNP thresholds were chosen using both the novel mixture distribution method implemented in TRACS and the Youden index as used previously to consider the person-to-person transmission landscape of the gut and oral microbiomes (4) (Supplementary Figure 3). No strain sharing is expected between different cohorts, and thus inferred transmissions between cohorts are highly likely to be false positives. In contrast, we expect to see bacterial strains persist within a patient’s gut microbiome over time, providing a valuable real-world ground truth dataset for method comparison. However, it is likely that strain persistence estimates will have a false positive at a rate similar to those observed in the between-cohort results.

**Fig. 4.**
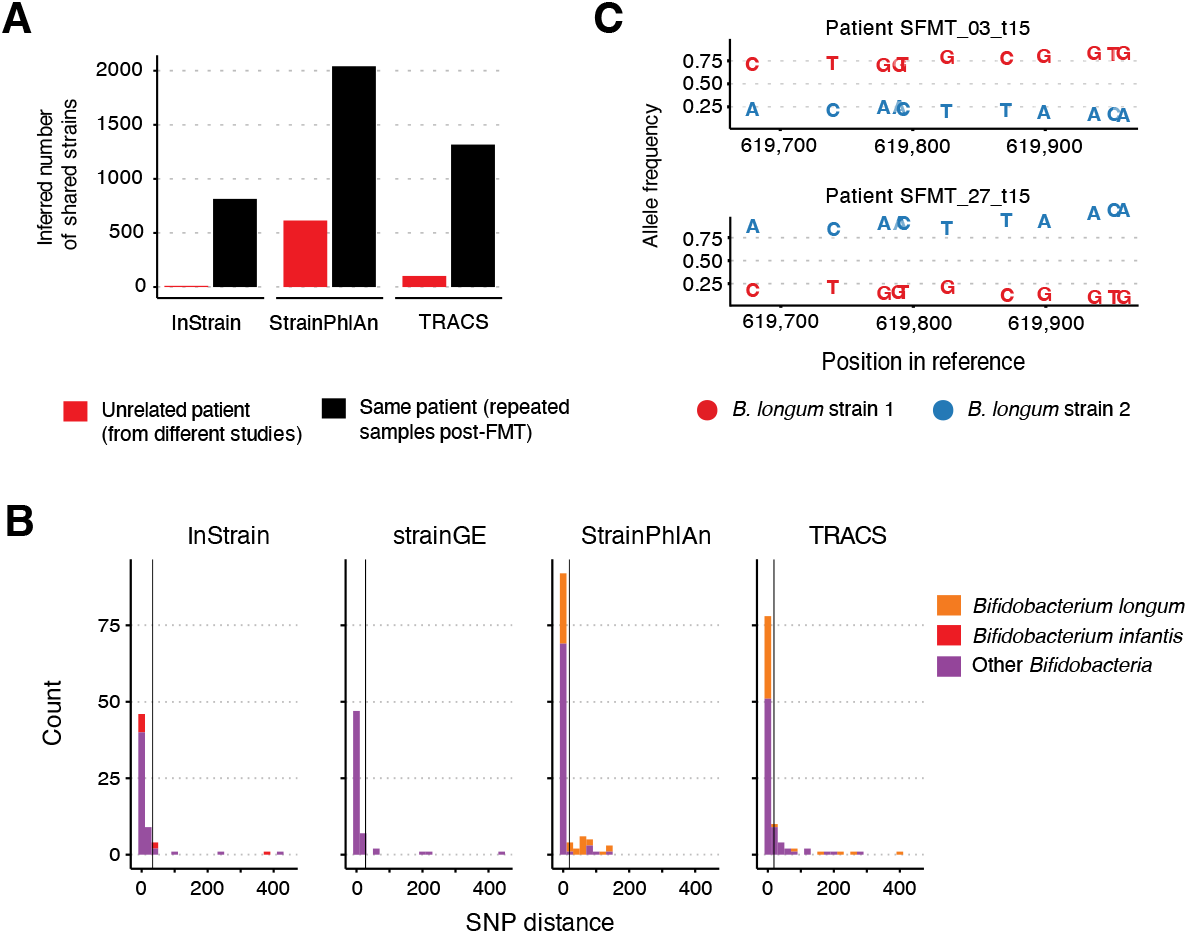
Estimates of engraftment in FMT triads. A) The inferred number of plausible transmission pairs (shared strains), inferred by each algorithm, between samples taken as part of study investigating the impact of Faecal Microbiota Transplants (FMTs). Transmission between samples from unrelated participants (shown in red) are highly likely to be false positives. In contrast, samples from the same participant following FMT (shown in black) are more likely to be true positives, with an error rate proportional to that observed in the unrelated participant results. Species specific SNP thresholds for identifying recent transmission were inferred using the mixture distribution approach (see Methods). An analogous plot using the Youden method as described in Valles-Colomer et al.(4), is given in Supplementary Figure 3 B) Histograms indicating the distribution of pairwise SNP distances (truncated at 500 SNPs) between Donor and recipient (post-transplant) samples for major Bifidobacterial species. Vertical grey lines indicate the SNP thresholds inferred using the TRACS mixture distribution method. InStrain identified the transmission of *B. infantis*, which is likely *B. longum* but mislabeled in the UHGG genome collection database, whereas StrainGE identified no transmission of *B. longum*. C) An example of multiple strains of *B. longum* being transmitted between a single donor and multiple recipients was detected exclusively by TRACS. The allele frequencies at polymorphic loci within a segment of the *B. longum* reference genome are shown for each recipient sample, with colours indicating the two distinct strains. The red strain is dominant in recipient sample SFMT_03_115 but in the minority in a separate recipient (SFMT_27_115) who received the same donor stool.

StrainPhlAn had the highest false positive rate which increased substantially using the Youden method for selecting SNP thresholds (Figure 4A, Supplementary Figures 3 & 4). Consistent with our simulation results, InStrain was the most conservative method, performing well when there is high sequence similarity between the reference and transmitted genomes. However, InStrain frequently overestimated SNP distances in cases of greater sequence divergence from the reference or when multiple strains of the same species were involved, as indicated by our previous simulation based analysis. In contrast to alternative algorithms, TRACS achieved a high sensitivity, with a low false positive rate.

To investigate this further, we considered the *Bifidobacteria* genus, and in particular the transmission of *Bifidobacterium longum*, a common pioneer coloniser of the infant gut microbiota(35, 36) which is often used as an infant probiotic. Understanding strain persistence and transmission is crucial for developing effective microbiome-based therapeutics. InStrain and StrainPhlAn were run using their default databases, while strainGE was supplied with a representative set of *Bifidobacteria* reference genomes following the user guide (Methods). TRACS was run using the GTDB reference database(37).

TRACS was the most sensitive algorithm at detecting the transmission of *B. longum*, identifying 31 shared strains, six more than the next most sensitive method, StrainPhlAn, which identified 25 (Figure 4B). InStrain and StrainGE proved to be the most conservative algorithms, while Strain-PhlAn identified the highest rate of sharing. This high rate of sharing likely includes many false positives due to Strain-PhlAn’s reliance on a small set of representative genes. Despite having *B. longum* genomes present in the reference database, StrainGE failed to identify this species amongst the engrafted strains. In contrast, InStrain identified 6 *B. infantis* transplanted strains as *B. longum* and *B. infantis* are considered as one species in InStrain’s default database (UHGG). Unlike competing algorithms, TRACS consistently identified cases of engraftment involving multiple strains of *B. longum* present within a single sample. One example is given in Figure 4C, where the frequency of the engrafted strains was reversed in two patients who received the same donor stool. TRACS was the only algorithm to correctly identify that these patients shared similar strains. Identifying cryptic transmission in studies such as these is crucial for developing accurate models of microbiome colonisation dynamics, a key component of efforts to identify strains with health benefits to develop microbiome-based therapies.

### Characterisation of maternal strain transmission and persistence across a large UK infant birth cohort

The transmission and subsequent colonisation events that drive the development of our gut microbiota can have important implications on our health in childhood and later life. In particular, the impact of perturbations driven by interventions in early childhood, such as caesarean section and antimicrobial treatment, have been associated with health complications including asthma and atopy (38, 39).

To characterise the transmission and persistence of bacteria at the strain level during early childhood, we analysed faecal samples from 1,288 healthy, full-term infants, sequenced as part of the UK BabyBiome Study(35, 36). Faecal samples were collected from all babies at least once during the neonatal period (≤ 1 month), with subsequent sampling from 302 infants, including 29 twins and one triplet, during later infancy (8.75 ± 1.98 months). Maternal faecal samples were also taken from 175 mothers, corresponding to 178 neonates. The TRACS algorithm was run on all samples using the GTDB reference database and SNP thresholds were calculated to distinguish recent transmission at the species level using the mixture distribution approach (as described in Methods). Consistent with the low error rate of TRACS and stringent infection control procedures in each hospital, the occurrence of shared strains between unrelated children, both within the same hospital and across different hospitals, was very low (0.78% and 0.73%, respectively; Figure 5A). Interestingly, the highest rate of strain sharing occurred between siblings, reflecting their closely shared strain colonisation history (from a common reservoir or between each other). Infants born via caesarean section exhibited markedly reduced strain sharing compared to those delivered vaginally (Figure 5B), aligning with findings from previous studies (35, 36, 40). *Bifidobacterium bifidum, Bifidobacterium longum, Phocaeicola vulgatus, Parabacterioides disasonis* and *E. coli* showed some of the largest differences in transmission rates between mothers and infants born vaginally compared to those born via caesarean section.

**Fig. 5.**
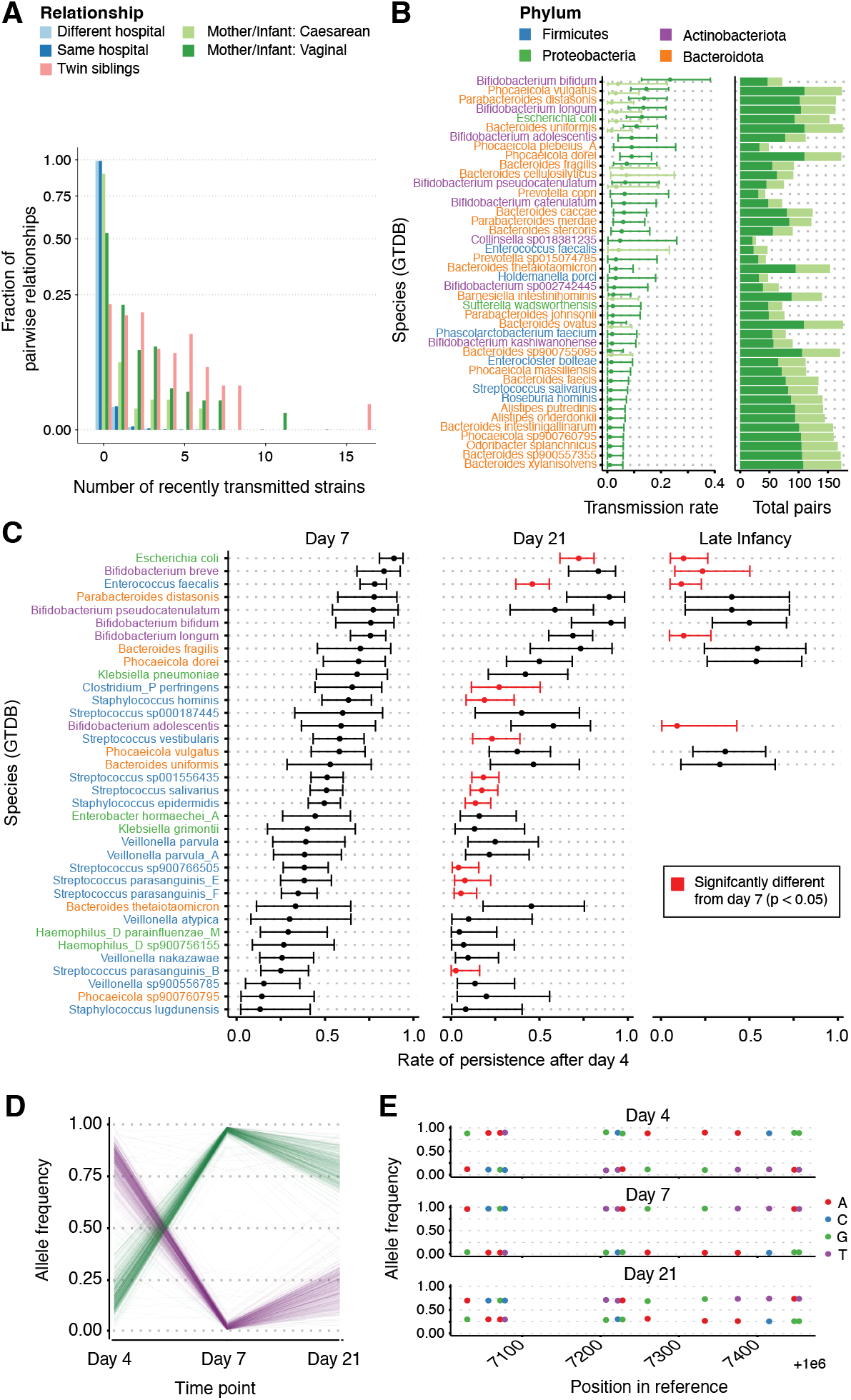
Maternal strain transmission and persistence across a large UK infant birth cohort. A) A bar plot indicating the fraction of pairwise relationships that involve a putative recent transmission versus the number of shared strains. SNP distances and species-specific thresholds are inferred using the TRACS algorithm. Relationships between hosts are represented by different colours. The low false positive rate of transmission is evident, as the vast majority of inter-hospital relationships involve zero transmissions. B) Species-specific transmission rates between mothers and babies. Point estimates and error bars are inferred using a binomial model of transmission if a mother is colonised (see Supplementary Methods). Points are coloured by the mode of delivery (as in A), highlighting the species-specific impact of delivery. The total number of possible transmission pairs (mothers colonised) is shown in the accompanying horizontal bar plot. C) The rate of strain persistence by species from birth to 7 days, 21 days, and late infancy. Persistence rates that differ significantly from day 7 are shown in red (Chi-squared test). Estimates are displayed only when at least 20 babies, who were originally observed to carry a species, are sampled again at respective time points. Point estimates and error bars are inferred using a binomial model. D) An example demonstrating the persistence of multiple distinct strains of *B. breve* in a single infant (ID: B00560) is presented. This example was not identified by StrainPhlAn, which only considers the dominant genotype. The frequencies of 1000 randomly chosen alleles, found to be at intermediate frequency on day 4, are shown. Initially, the purple strain dominates, but by days 7 and 21, the green strain becomes dominant. E) Plotting of individual allele frequencies of two *B. breve* strains (from panel D) relative to a portion of the *B. breve* reference genome.

In addition to tracking strain transmission between hosts, TRACS can monitor the longitudinal persistence of strains within a single host. This includes observing the strain stability in infants, which was found to vary significantly between species (Figure 5C). Strains of commonly pathogenic species such as *E. coli* and *E. faecalis* were maintained for relatively short periods of time and are generally eliminated or replaced by late infancy. The persistence of the pioneer colonisers in the neonatal gut, including *B. longum* and *B. breve* strains(36), remained high during the first three weeks of infancy but declined markedly in late infancy, likely due to changes in diet and decreased breastfeeding rates (41, 42). In contrast, some other maternally transmitted commensal genera such as *B. bifidum* and closely related Phocaeicola, Bacteroides and Parabacteroides species were maintained at higher rates for the entire duration of the study. Critically, the TRACS algorithm identified 40.5% (49/121) additional strain sharing events compared to StrainPhlAn. This indicates instances of repeated co-colonisation of infants with multiple strains of *B. breve*, which were overlooked by algorithms like StrainPhlAn that consider only the dominant genotype (Supplementary Figure 6). For instance, Figure 5D and 5E illustrate the persistence of two *B. breve* strains over three time points within a single infant. Initially, the purple strain is dominant with a mean allele frequency of 0.82 (±0.081) but then drops to lower frequencies of 0.063 (±0.181) and 0.249 (±0.156) respectively at later time points. Taken together, these results demonstrate the effectiveness of the TRACS algorithm in analysing the fine-scale transmission and persistence of strains within the human microbiota.

## Discussion

TRACS is a versatile and modular algorithm for inferring recent pairwise transmission events from metagenomic and single species population sequencing data. The algorithm addresses several key sources of error, including sequencing errors, uneven sequencing coverage, shared sequence homology between microbes. Importantly, TRACS also accounts for the presence of multiple strains of the same species within a sample. Using simulations and artificial laboratory mixtures of *S. pneumoniae*, we show that accounting for shared homology and recombination substantially improves the accuracy of TRACS compared to alternative algorithms.

The scalability and versatility of the TRACS algorithm allows it to be applied to a broad spectrum of pathogens including viruses, bacteria and parasites. This includes contexts where species specific whole genome bioinformatics transmission pipelines are yet to be developed. For example, although whole genome sequencing is not typically used to track the transmission of malaria-causing parasites, we demonstrate using single-cell genomics, that by omitting the complex and error-prone step of deconvolution, the TRACS algorithm effectively identifies shared strains of *P. falciparum* between samples. By estimating a lower bound on the SNP distance between strains in two samples, TRACS can reliably exclude recent transmission events. Combined with its versatility, computational efficiency, and ability to incorporate sampling dates through an enhanced version of the TransCluster model, TRACS is highly suitable for use in routine public health surveillance pipelines.

Beyond pathogen surveillance, TRACS can track the colonisation and carriage dynamics of the human microbiota. Using well-characterised metagenomic datasets from FMT studies, we demonstrate that the TRACS algorithm, which includes a novel method for estimating transmission distance thresholds, offers a superior balance of sensitivity and specificity for tracking strain acquisition and persistence within the gut. Notably, TRACS identifies instances of minority strain transmission between FMT donors and recipients, as well as longitudinally sampled infants, which are not detected by alternative tools. Moreover, in addition to confirming that caesarean sections significantly reduce mother-to-baby bacterial transmission, we identified species-specific differences in colonisation and persistence rates. Notably, *E. faecalis* and *B. cellulosilyticus* had higher transmission rates following caesarean delivery. As with other genomic transmission studies, it is essential to consider the broader epidemiology of host interactions. For example, in some cases, both mother and infant may be independently colonised by the same strain from the hospital environment or other unsampled sources, such as family members.

TRACS was developed to identify recent transmission events, enhancing both accuracy and ease of use. However, the algorithm was not developed for identifying high-quality within-host variants, making it unsuitable for detecting selection through dN/dS ratios. Furthermore, since the algorithm provides only a lower bound estimate of transmission or SNP distance, it cannot accurately infer genetic relationships over thousands of SNPs that represent much longer evolutionary timescales. For these purposes, tools like inStrain and Strain-PhlAn respectively, are more appropriate. Similar to most minority variant–calling algorithms, TRACS requires sufficient sequencing depth to distinguish true variants from sequencing errors. Consequently, by default, TRACS requires a minimum coverage of 5x coverage to accurately estimate distances between strains. As with other metagenomic transmission inference methods, TRACS does not determine transmission direction. For this, we recommend using TRACS to identify potential transmission events, followed by more computationally intensive phylogenetic methods that incorporate additional epidemiological information (8, 43).

Metagenomics and deep population sequencing present a powerful alternative to relying solely on representative genomes for tracking the transmission and persistence of both pathogenic and commensal microbes. The versatility and robustness of the TRACS algorithm allows for the simultaneous analysis of multiple strains and species, which will help advance our understanding of how microbes transmit, establish, and persist within their hosts.

## Methods

### TRACS algorithm

The TRACS algorithm consists of three central modules that can be used independently (see Figure 1). The **alignment** module generates a polymorphism-aware alignment against each reference genome found within a given sample. Given either a multi-sample reference based alignment or a traditional Multiple Sequence Alignment (MSA), the **distance** module estimates pairwise SNP distances. Optionally, the distance module can also estimate the expected number of intermediate hosts separating pairs of samples using an extended version of the TransCluster algorithm(17). The TransCluster algorithm requires estimates of the transmission generation time (transmissions/year) and clock rate (SNPs/genome/year) of the species to be provided. Finally, the **cluster** module estimates putative transmission clusters using single-linkage hierarchical clustering.

### Alignment

The alignment module takes raw sequence data as input and produces alignments to multiple reference genomes in FASTA format. IUPAC ambiguity codes are used to represent polymorphic sites in each alignment, enabling the estimation of SNP distances between populations. The alignment module supports a variety of database formats tailored to specific applications. Users can provide a single reference, utilise a Sourmash database that incorporates Genbank genome IDs, or opt for a custom TRACS database. The latter includes both reference genomes and a corresponding Sourmash database, which can be constructed using the TRACS build-db command.

If a database containing multiple references is provided, the algorithm will first select those represented in a sample using the Sourmash gather command (14). Minimap2 (44) is then used to align all reads within a sample to each reference independently and a pileup of alleles identified at each site in the reference is generated using htsbox (45).

To control for variable sequencing coverage, we developed an empirical Bayes method. This is applied to each genome alignment independently. The counts of minority variants at each site within a single reference alignment are modelled jointly using a multinomial Dirichlet distribution. This allows for the estimation of the posterior frequency of variants across an individual alignment by considering the average frequency of minority variants at polymorphic sites as follows. Assume *n*_*i*1_ is the read count of the most frequent allele at polymorphic site *i* in an alignment with *n*_*i*2_, *n*_*i*3_ and *n*_*i*4_ being the counts of the 2nd, 3rd and 4th most frequent allele respectively. Our goal is to estimate a prior on the expected frequency of each allele at a given site by considering the observed distribution of allele frequencies across all sites in the dataset.

We assume a Dirichlet prior on the joint frequency (**p**_**i**_ = (*p*_*i*1_, *p*_*i*2_, *p*_*i*3_, *p*_*i*4_)) of the counts *n*_*i*_, with parameter vector *–* such that the probability density can be written as:

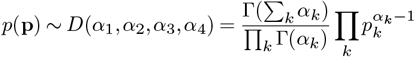

Assuming the resulting read counts at each site are drawn from a multinomial distribution, the marginal likelihood is the Dirichlet-multinomial distribution given by:

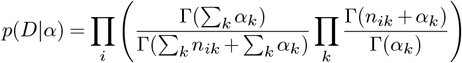

where *D* indicates the vector of read counts at each site *i*: *D* = {**x**_1_, …, **x**_*N*_} where **x**_*i*_ = {*n*_*i*1_, *n*_*i*2_, *n*_*i*3_, *n*_*i*4_}

Point estimates of the *–* parameters were made by maximising the marginal likelihood using the fixed point iteration method (46). The maximum a posteriori estimate of the frequency of each allele under this prior at each site can then be estimated as:

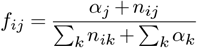

After estimating posterior frequencies at all variable sites in an alignment, TRACS applies a frequency cut-off which requires a coverage of at least 5 reads by default. Sites falling below this threshold are then excluded from further consideration in subsequent estimates of SNP distances. Furthermore, to control for major outliers in sequencing coverage, regions with coverage exceeding 1.5 times the interquartile range above or below the quartiles, as identified by Tukey’s method, are excluded (47).

### Distance

Pairwise SNP distances are estimated using a bitset-based algorithm to enhance both the speed and memory efficiency of the process, whilst incorporating IUPAC ambiguity codes. Each sequence in a Multiple Sequence Alignment is represented as a matrix of bits (0’s and 1’s), where rows indicate the four possible alleles and columns indicate the position in the sequence. Sites where multiple different alleles are observed are then represented by multiple 1’s within a single column. A series of bitwise AND operations are used per allele, and the results are combined with OR operations to rapidly calculate the sites which share common alleles between each pair of samples.

To account for the impacts of recombination and shared homology between strains, we include an optional additional step to identify regions within the pairwise comparison that have an elevated rate of SNPs. This uses a similar scan statistic to that used in popular recombination aware phylogenetic reconstruction algorithms (16). While this step may exclude sites that could be informative for transmission, SNPs introduced by recombination are rarely incorporated into public health genomic pipelines due to the challenges they pose for phylodynamic analyses. Additionally, the noise introduced by cross-mapping from shared homology is likely to outweigh any potential benefits of retaining these sites.

Initially, the rate of SNPs within the alignment is calculated as 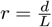, where *d* is the total number of SNPs and *L* is the alignment length. For each pairwise comparison, a window size *w*_*l*_ is chosen such that the expected number of SNPs within the window is equal to one: *E*(*n*| *p, w*_*l*_)= 1. The maximum window size is set to 10kb. At each SNP location, *i*, the number of SNPs, *s*_*i*_, within the window centred at *i* is calculated. Assuming a binomial distribution, the probability of observing at least *s*_*i*_ SNPs is given by:

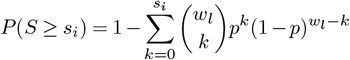

A p-value threshold is then specified to exclude SNPs that are centred within a window that has a significantly high rate of SNPs (*p* = 0.05 by default). A Bonferroni correction is made to control for multiple testing (48). For SNPs that fall within a distance *w*_*l*_*/*2 of either edge of the alignment, *w*_*l*_ is truncated.

### Modified TransCluster algorithm

While SNP distances can be useful in distinguishing recent transmission events, they do not incorporate the sampling times of the hosts being considered. To incorporate the sampling times, TRACS includes an extended version of the TransCluster algorithm that both improves its speed and allows for the estimation of the expected number of intermediate hosts separating two samples.

Assuming SNPs accumulate at rate *γ* and an average epidemiological generation time (the time between transmission events) *β*, the TransCluster algorithm can be used to estimate the probability of *k* intermediate hosts occuring along the transmission chain between a pair of sampled hosts (17). Given a SNP distance of *N* and a time separation of *δ* between the two samples, Stimson et al.(17) showed that the distribution of *k* intermediate hosts could be written as:

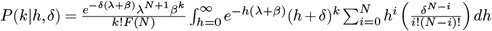

where *h* is the evolutionary time separating two samples. In the original implementation of TransCluster, this expression was computed via numerical integration. This method can become computationally prohibitive when considering thousands of pairwise comparisons. We show that this equation can be re-written as a finite sum that can be solved rapidly using efficient caching of intermediate values:

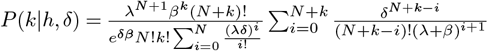

A full derivation is given in the supplementary methods. Given this derivation, the expected number of intermediate hosts can be written as:

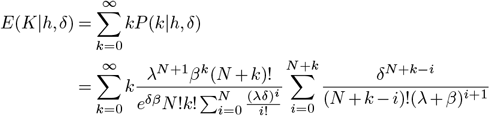

While this is an infinite sum, an upper bound on the error after truncating at a given *k* can be calculated (supplementary methods) allowing for the sum to be calculated to a user specified precision. Again, TRACS makes use of efficient caching strategies to reduce the number of times this equation is calculated.

### Clustering

Consistent with other transmission inference pipelines, TRACS implements single linkage hierarchical clustering to identify putative transmission clusters. The **cluster** module takes a list of pairwise distance estimates output from the **distance** module and a user supplied SNP or transmission distance threshold and outputs a CSV file indicating which genome belongs to each cluster. Guidance on the selection of an appropriate threshold is given below. Single linkage clustering is preferable as it allows for long transmission chains to be grouped into a single cluster.

### Inferring a transmission threshold

To determine an appropriate SNP threshold for distinguishing recent transmission, we assume that it is possible to provide two sets of samples where the first includes possible transmission events and the second is unlikely to include any recent transmission. By fitting a mixture distribution to these samples, we can calculate a suitable threshold.

We assume that SNP distances between pairs of samples that are unlikely to be related by recent transmission (*d*_*u*_) are distributed according to a Negative Binomial distribution with parameters *n* and *p* such that

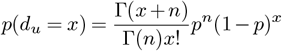

Moreover, we assume that the distribution of SNP distances between pairs of SNPs related by recent transmission (*d*_*r*_) follows a Poisson distribution with meanλ

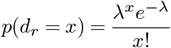

Thus the distribution of SNP distances in the dataset that contains both recent transmission and more distant transmission (*d*_*m*_) can be modelled as a mixture of these two distributions where *q* indicates the probability that any given pair of samples are related by recent transmission.

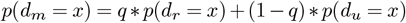

To estimate these parameters we fit the Negative Binomial model to the dataset containing exclusively distantly related samples using maximum likelihood. This produces a distribution of the SNP distance between distantly related samples based on the second dataset. Substituting these estimates into the mixture then allows for the estimation of the remaining parameters using maximum likelihood. Note that during this second estimation, we use the first data set without the second dataset. Finally a SNP threshold can be chosen by selecting the probability that a close transmission pair is erroneously excluded.

To select thresholds for the analysis of transmission in the mother and baby dataset(35, 36), repeated samples of the same baby were used as the ‘close transmission’ dataset and pairs of samples from different infants were used as the ‘distant’ dataset. In the FMT dataset, the ‘close transmission’ dataset consisted of samples from the same patient post-FMT and the ‘distant’ dataset consisted of pairs of samples taken from unrelated donors. In both cases, a probability of missing a transmission of 0.001 was chosen. To adjust for the fact that the ‘close transmission’ pairs were from the same patient, the estimated thresholds were multiplied by 3 to allow for twice the evolutionary time to occur in the mother/donor.

### Alternative algorithms and databases

In all cases, StrainPhlAn was run using version 4.0.5 with the default database using the command line parameters outlined in Valles-Colomer et al(4).

For the metagenomic simulations, TRACS, InStrain and StrainGE were run using a reference database comprising of the GTDB represenative genomes that correspond to the 11 bacterial species identified in a study investigating the transmission of drug-resistant pathogens using metagenomics(23).

For the pneumococcal mixed strain simulations, TRACS and StrainGE were run using representative genomes from each sequencing cluster described in the Global Pneumococcal Sequencing project (28). As InStrain required a dereplicated dataset, the 19F serotype included in the laboratory mixtures was used to run InStrain on the simulated pneumococcal deep population sequencing data.

The default databases for each tool were used in the analysis of the FMT metagenomic data(34). The GTDB r207 bacterial reference database was used (37) to run TRACS. InStrain was run using the UHGG reference database as described in the InStrain user manual. As it was computationally expensive to consider all possible species when running StrainGE on the FMT data, we restricted all analyses to the Bifidobacterium genus. Reference genomes were downloaded from Refseq and the database was constructed following the StrainGE user manual.

TRACS used the same pneumococcal reference database to analyse the pneumococcal deep sequencing data from the Maela refugee camp. To analyse the SARS-CoV-2 and *P. falciparum* deep population sequencing data, the reference genome from each species was used when running TRACS. Similar to the FMT data, TRACS was run using the GTDB r207 bacterial reference database to analyse the UK infant birth cohort.

To compare the resource requirements of each algorithm, we measured CPU time required for all simulated metagenomic samples where one genome was simulated to have been transmitted (Supplementary Figure 7). Memory was compared on a representative pair of samples. Comparing the results across algorithms is challenging, as each requires different tasks to be performed separately by the user, such as read alignment and SNP distance calculation. To ensure a fair comparison, we used the same tools employed by TRACS for operations not automatically handled by other pipelines. Although the exact differences in speed and memory usage may vary depending on the dataset, TRACS consistently demonstrated the lowest memory and CPU usage in this test, outperforming the next best algorithm by approximately a factor of 2 (Supplementary Figure 7).

The exact commands used to run StrainPhlAn, inStrain and StrainGE are given in the supplementary code provided in the associated GitHub repository (https://github.com/gtonkinhill/tracs_manuscript).

### Simulated datasets

Given a user-specified SNP distance and a set of reference genomes, transmission of an individual strain between pairs of metagenomic or population sequencing samples were simulated using a custom python script provided as part of the TRACS package.

The number of reference genomes included in each sample was simulated using a Poisson distribution. The user specified number of SNPs was then simulated by randomly introducing mutations along one reference genome. This genome was included in both samples in a pair. The remaining reference genomes in each sample were randomly selected from the user provided list. In cases where both samples shared a reference genome other than the transmitted one, additional SNPs were randomly introduced. The total number of SNPs separating non-transmitted strains was drawn from a Poisson distribution with mean 10,000. This value was chosen to ensure that all methods should be able to reliably distinguish the small genetic distances characteristic of recent transmission from much larger distances representing hundreds of years of evolution. While closely related strains can be shared through routes other than transmission, such events cannot be distinguished solely by genetic distance. As a result, they are not considered in detail in these simulations.

Simulated sequencing depths were drawn from a Dirichlet distribution with all parameters equal to 1. A cumulative read depth across all genomes of 500 reads was used and Illumina reads were generated using the ART synthetic read simulator(49), similar to the simulation method used in the development of StrainGE(10). This coverage corresponded to an average of 18 million and 7 million reads in the metagenomic and pneumococcal simulations, respectively. Simulated Nanopore reads were generated using Badread v0.4.1 (50). A full set of parameters and commands can be found in the supplementary code provided in the GitHub repository. To simulate transmission between gut metagenomes, 11 bacterial species were selected from a study investigating the transmission of drug-resistant pathogens using metagenomics(23). A series of high quality reference genomes were then selected from the full GTDB r207 database that averaged at 100%, 99% and 97% in pairwise sequence identity from the GTDB r207 representative genomes for each species. Simulated transmission pairs were then generated at each identity level. To increase the complexity of the metagenomic simulations, and inline with previous simulation approaches(9, 10), real metagenomic samples were appended to the in-silico simulated data. A pair of samples were chosen from the FMT data set for this purpose which were found not to share strains by all algorithms (accessions ERR9707885 and ERR9709032).

To simulate transmission among deeply sequenced populations of pneumococci, we randomly selected 28 pneumococcal genomes from a prior study (51). A full list of all commands used along with the reference genomes is provided in the associated GitHub repository.

## Data availability

SARS-CoV-2 sequencing reads were taken from a recent study investigating the within-host evolution of the virus(26). The ENA IDs of the 37 SARS-CoV-2 samples found to contain multiple distinct lineages are listed in the supplementary GitHub repository. All sequencing reads from the original study are available in the ENA under accession number ERP126512. The *S. pneumoniae* and *P. falciparum* sequencing reads are available from the ENA under project codes PRJEB22771 and PRJNA482776 respectively(7, 33). The deep metagenomic sequencing reads of gut samples from mothers and their children is available from the ENA under the accession number ERP115334(35, 36). The FMT triad samples included sequencing data published in the ENA under project code PRJEB47909(34). The sample accession codes used in all analyses are provided in the supplementary files on GitHub.

## Code availability

The TRACS code and user documentation is available under an MIT open source licence on GitHub at https://github.com/gtonkinhill/tracs/. Supplementary code used to generate the analysis and figures in this manuscript is available at https://github.com/gtonkinhill/tracs_manuscript.

