## Supplementary methods for "Enhanced metagenomics-enabled transmission inference with TRACS"

### TRACS Supplementary Methods

August 19, 2024

#### 1 TransCluster model

The TransCluster algorithm improves on a simple SNP distance metric by accounting for the time separating two samples [1]. It assumes both the substitution rate and mean generation time are known. We provide a brief description of this approach as well as some extensions that improve its scalability and interpretability.

Let $N$ be the SNP distance separating two genomes, $\delta$ the time difference between when the samples were taken, and $h/2$ the time between the earlier sample and their MRCA. We would like to estimate $h + \delta$, the total time along both lines of descent between the MRCA and the two sample times.

The number of SNPs per unit time can be modelled as a Poisson process with evolutionary rate $\lambda$. We first decompose $N$ such that $N = N_h + N_\delta$ where $N_h$ represents the number of SNPs that accumulated before sampling the first case and $N_\delta$ accounts for any SNPs that accumulated in the subsequently sampled case after the initial sample has been taken (see Figure 1).

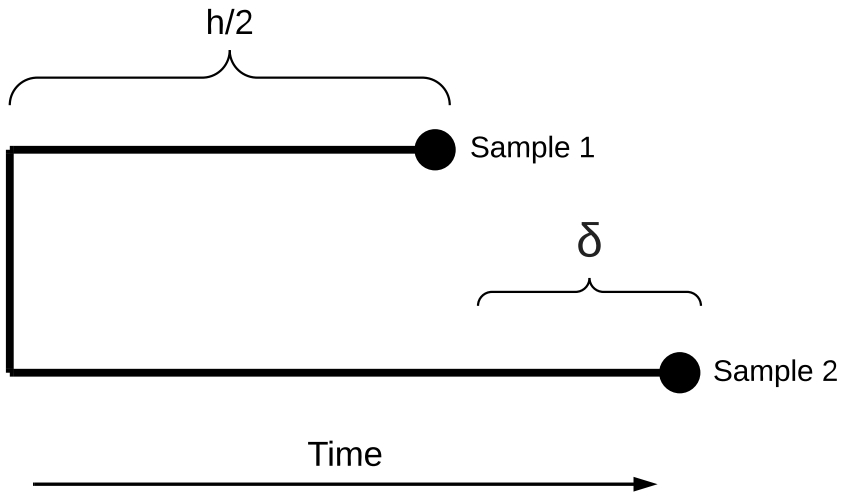


Figure 1: A diagram describing the parameters used in the Transcluster algorithm. $\delta$ indicates the time between samples while $h$ indicates the total time along both lines of descent from the MRCA to the time the first sample was taken. Thus, the time between the MRCA and the first sample is $h/2$.

Without this restriction $N_\delta$ would be Poisson distributed with parameter $\lambda\delta$. After conditioning on the fact that the number of SNPs $N_\delta$ can not be greater than the total number of SNPs observed ($0 \leq N_\delta \leq N$) we get the distribution of $N_\delta$ as

$$P(N_\delta|\delta) = \left(\frac{e^{-\lambda\delta}(\lambda\delta)^{N_\delta}}{F(N_\delta)}\right) \tag{1}$$

where $F$ is the CDF, $F(N_\delta) = \sum_{i=0}^{N_\delta} e^{-\lambda\delta}(\lambda\delta)^i/i!$

We now consider $N_h$, the number of SNPs accumulated prior to sampling. This can be modelled as a Poisson process with rate $\lambda$ and thus the waiting time until the $N_h$th SNP as described in Stimson *et al.*, [1], can be given by

$$\frac{\lambda^{N_h} h^{N_h-1} e^{-\lambda h}}{(N_h-1)!} \tag{2}$$

As we know that $N_h$ SNPs have already accumulated we are interested in the waiting time for the $(N_h+1)$th SNP. Thus we can write the likelihood of $h$ given $N_h$ as

$$\mathcal{L}(h|N_h) = \frac{\lambda^{N_h+1} h^{N_h} e^{-\lambda h}}{N_h!} \tag{3}$$

To find the likelihood $h$ given $N$ and $\delta$ we sum over all possible values of $N_\delta$ as

$$\mathcal{L}(h|N,\delta) = \sum_{i=0}^{N} \mathcal{L}(h|i,\delta) P((N-i)|\delta) \tag{4}$$

which can be expanded as

$$\mathcal{L}(h|N,\delta) = \frac{e^{-\lambda(h+\delta)}\lambda^{N+1}}{F(N)} \sum_{i=0}^{N} \frac{h^i \delta^{N-i}}{i!(N-i)!} \tag{5}$$

Given $h$ and $\delta$, we now assume that the generation time (the time between transmission events) is parameterised by $\beta$, the rate at which the pathogen jumps to a new host. We assume $\beta$ is constant resulting in another Poisson process for the number of intermediate hosts between the two samples. This can be expressed as

$$P(k|h,\delta) = \frac{\beta^k (h+\delta)^k e^{\beta(h+\delta)}}{k!} \tag{6}$$

integrating over $h$ gives

$$P(k|h,\delta) = \int_{h=0}^{\infty} \mathcal{L}(h|N,\delta) P(k|h) dh \tag{7}$$

this expression can be expanded as described in [1], to give

$$P(k|h,\delta) = \frac{e^{-\delta(\lambda+\beta)}\lambda^{N+1}\beta^k}{k!F(N)} \int_{h=0}^{\infty} e^{-h(\lambda+\beta)}(h+\delta)^k \sum_{i=0}^{N} h^i \left(\frac{\delta^{N-i}}{i!(N-i)!}\right) dh \tag{8}$$

#### 1.1 A fast solution to Equation 8

In the original implementation of Transcluster, Equation 8 is solved via numerical integration. Whilst this is suitable for a small number of comparisons when dealing with a very large number of genomes it can become a bottleneck in analysis. Instead the integral can be expressed as a finite sum as

$$
\begin{aligned}
&\int_{h=0}^{\infty} e^{-h(\lambda+\beta)}(h+\delta)^k \sum_{i=0}^{N} h^i \left(\frac{\delta^{N-i}}{i!(N-i)!}\right) dh \\
&= \int_{h=0}^{\infty} e^{-h(\lambda+\beta)}(h+\delta)^k \sum_{i=0}^{N} \left(\frac{h}{\delta}\right)^i \frac{\delta^N}{N!}\binom{N}{i} dh \\
&= \frac{1}{N!}\int e^{-h(\lambda+\beta)}\,(h+\delta)^k \left(1+\frac{h}{\delta}\right)^N dh \\
&= \frac{1}{N!}\int e^{-h(\lambda+\beta)}\,(h+\delta)^{N+k}\,dh \\
&= \frac{1}{N!}\int e^{-h(\lambda+\beta)} \sum_{i=0}^{N+k} h^i \delta^{N+k-i}\binom{N+k}{i} dh \\
&= \frac{1}{N!}\sum_{i=0}^{N+k} \delta^{N+k-i}\binom{N+k}{i}\int e^{-h(\lambda+\beta)} h^i dh \\
&= \frac{1}{N!}\sum_{i=0}^{N+k} \binom{N+k}{i}\delta^{N+k-i}\frac{\Gamma(i+1)}{(\lambda+\beta)^{i+1}}
\end{aligned}
\tag{9}
$$

Substituting this term into Equation 8 and simplifying a bit gives

$$
P(k|h,\delta) = \frac{\lambda^{N+1}\beta^k (N+k)!}{e^{\delta\beta} N!k! \sum_{i=0}^{N}\frac{(\lambda\delta)^i}{i!}} \sum_{i=0}^{N+k} \frac{\delta^{N+k-i}}{(N+k-i)!(\lambda+\beta)^{i+1}} \tag{10}
$$

#### 1.2 Expected $K$

To identify potential transmission links a threshold on the probability of a transmission occurring within $K$ intermediate hosts is usually chosen. i.e. ($P(k \leq K) <$ threshold). We have found that some users find selecting both a suitable K and the threshold to be confusing and the resulting probabilities can be harder to interpret. Instead, estimating the $E(K)$ can provide a more readily interpretable distance.

$$
\begin{aligned}
E(K|h,\delta) &= \sum_{k=0}^{\infty} kP(k|h,\delta) \\
&= \sum_{k=0}^{\infty} k \frac{\lambda^{N+1}\beta^k (N+k)!}{e^{\delta\beta} N!k! \sum_{i=0}^{N}\frac{(\lambda\delta)^i}{i!}} \sum_{i=0}^{N+k} \frac{\delta^{N+k-i}}{(N+k-i)!(\lambda+\beta)^{i+1}}
\end{aligned}
\tag{11}
$$

Now, if we assume $\delta$, $\lambda$ and $\beta$ are all positive we have

$$
\begin{aligned}
&\sum_{i=0}^{N+k} \frac{\delta^{N+k-i}}{(N+k-i)!(\lambda+\beta)^{i+1}} \\
&= \frac{1}{(\lambda+\beta)^{N+k+1}} \sum_{i=0}^{N+k} \frac{(\delta(\lambda+\beta))^{N+k-i}}{(N+k-i)!} \\
&= \frac{1}{(\lambda+\beta)^{N+k+1}} \sum_{m=0}^{N+k} \frac{(\delta(\lambda+\beta))^{m}}{m!} \\
&< \frac{1}{(\lambda+\beta)^{N+k+1}} \sum_{m=0}^{\infty} \frac{(\delta(\lambda+\beta))^{m}}{m!} \\
&= \frac{e^{\delta(\lambda+\beta)}}{(\lambda+\beta)^{N+k+1}}
\end{aligned} \tag{12}
$$

Substituting this into $E(K)$ allows us to construct an upper bound on the error remaining if we truncate the series in equation 11 (The truncated series is already a lower bound). We find the upper bound on the overall expected value as

$$
\begin{aligned}
E(K|h,\delta) &= \sum_{k=0}^{\infty} kP(k|h,\delta) \\
&< \sum_{k=0}^{\infty} k \frac{\lambda^{N+1}\beta^k (N+k)!}{e^{\delta\beta} N! k! \sum_{i=0}^{N} \frac{(\lambda\delta)^i}{i!}} \frac{e^{\delta(\lambda+\beta)}}{(\lambda+\beta)^{N+k+1}} \\
&= \frac{\lambda^{N+1} e^{\delta(\lambda+\beta)}}{e^{\delta\beta} N! \sum_{i=0}^{N} \frac{(\lambda\delta)^i}{i!}} \sum_{k=0}^{\infty} \frac{k\beta^k (N+k)!}{(\lambda+\beta)^{N+k+1} k!} \\
&= \frac{\beta e^{\delta\lambda}(N+1)}{\lambda \sum_{i=0}^{N} \frac{(\lambda\delta)^i}{i!}}
\end{aligned} \tag{13}
$$

Now,

$$
\begin{aligned}
E(K|h,\delta) &= \sum_{k=0}^{\infty} kP(k|h,\delta) \\
&= \sum_{k=0}^{M-1} kP(k|h,\delta) + O(M)
\end{aligned} \tag{14}
$$

where

$$
O(M) < \frac{\beta e^{\delta\lambda}(N+1)}{\lambda \sum_{i=0}^{N} \frac{(\lambda\delta)^i}{i!}} - \sum_{k=0}^{M} k \frac{\lambda^{N+1}\beta^k (N+k)!}{e^{\delta\beta} N! k! \sum_{i=0}^{N} \frac{(\lambda\delta)^i}{i!}} \frac{e^{\delta(\lambda+\beta)}}{(\lambda+\beta)^{N+k+1}} \tag{15}
$$
